# A phylogenomic resolution of the sea urchin tree of life

**DOI:** 10.1101/430595

**Authors:** Nicolás Mongiardino Koch, Simon E. Coppard, Harilaos A. Lessios, Derek E. G. Briggs, Rich Mooi, Greg W. Rouse

**Affiliations:** Department of Geology and Geophysics, Yale University, New Haven CT, USA; Department of Biology, Hamilton College, Clinton NY, USA; Smithsonian Tropical Research Institute, Balboa, Panama; Peabody Museum of Natural History, Yale University, New Haven CT, USA; Department of Invertebrate Zoology and Geology, California Academy of Sciences, San Francisco CA, USA; Scripps Institution of Oceanography, UC San Diego, La Jolla CA, USA

**Keywords:** Echinoidea, sea urchins, sand dollars, phylogenomics, genome, transcriptome

## Abstract

**Background:** Echinoidea is a clade of marine animals including sea urchins, heart urchins, sand dollars and sea biscuits. Found in benthic habitats across all latitudes, echinoids are key components of marine communities such as coral reefs and kelp forests. A little over 1,000 species inhabit the oceans today, a diversity that traces its roots back at least to the Permian. Although much effort has been devoted to elucidating the echinoid tree of life using a variety of morphological data, molecular attempts have relied on only a handful of genes. Both of these approaches have had limited success at resolving the deepest nodes of the tree, and their disagreement over the positions of a number of clades remains unresolved.

**Results:** We performed de novo sequencing and assembly of 17 transcriptomes to complement available genomic resources of sea urchins and produce the first phylogenomic analysis of the clade. Multiple methods of probabilistic inference recovered identical topologies, with virtually all nodes showing maximum support. In contrast, the coalescent-based method ASTRAL-II resolved one node differently, a result apparently driven by gene tree error induced by evolutionary rate heterogeneity. Regardless of the method employed, our phylogenetic structure deviates from the currently accepted classification of echinoids, with neither Acroechinoidea (all euechinoids except echinothurioids), nor Clypeasteroida (sand dollars and sea biscuits) being monophyletic as currently defined. We demonstrate the strength and distribution of phylogenetic signal throughout the genome for novel resolutions of these lineages and rule out systematic biases as possible explanations.

**Conclusions:** Our investigation substantially augments the molecular resources available for sea urchins, providing the first transcriptomes for many of its main lineages. Using this expanded genomic dataset, we resolve the position of several clades in agreement with early molecular analyses but in disagreement with morphological data. Our efforts settle multiple phylogenetic uncertainties, including the position of the enigmatic deep-sea echinothurioids and the identity of the sister clade to sand dollars. We offer a detailed assessment of evolutionary scenarios that could reconcile our findings with morphological evidence, opening up new lines of research into the development and evolutionary history of this ancient clade.

## Background

Echinoidea Leske, 1778 is a clade of marine animals including species commonly known as sea urchins, heart urchins, sand dollars and sea biscuits. It constitutes one of the five main clades of extant Echinodermata, typically pentaradially symmetric animals, which also includes highly distinctive components of the marine fauna such as sea lilies and feather stars (crinoids), starfish (asteroids), brittle stars (ophiuroids) and sea cucumbers (holothuroids). Fossil evidence suggests that these lineages, as well as a huge diversity of extinct relatives, trace their origins to the early Paleozoic [1, 2]. Their deep and rapid divergence from one another, coupled with long stem groups leading to the origin of extant forms, for a long time impeded a robust resolution of their interrelationships. Nonetheless, a consensus has emerged supporting a close relationship between echinoids and holothuroids, as well as between asteroids and ophiuroids, with crinoids as sister to them all [3–5].

A little over 1,000 extant species of echinoids have been described [6], comprising a radiation whose last common ancestor likely arose during the Permian [7, 8], although the stem of the group extends back to the Ordovician [9]. Extant echinoid species richness is vastly eclipsed by the more than 10,000 species that constitute the rich echinoid fossil record [10]. Nonetheless, it seems safe to assume that echinoid diversity at any given point in time has never exceeded that of the present day [11, 12]. Today, sea urchins are conspicuous occupants of the marine realm, inhabiting all benthic habitats from the poles to the Equator and from intertidal to abyssal zones [13]. As the main epifaunal grazers in many habitats, sea urchins contribute to the health and stability of key communities such as kelp forests [14] and coral reefs [15, 16]. Likewise, bioturbation associated with the feeding and burrowing activities of a large diversity of infaunal echinoids has a strong impact on the structure and function of marine sedimentary environments [17, 18]. Figure 1 provides a snapshot of their morphological diversity. Since the mid-19^th^ century, research on sea urchins has played a major role in modelling our understanding of animal fertilization and embryology [19, 20], with many species becoming model organisms in the field of developmental biology. This line of research was radically expanded recently through the application of massive sequencing methods, resulting in major breakthroughs in our understanding of the organization of deuterostome genomes and the gene regulatory networks that underlie embryogenesis [21, 22].

**Fig. 1.**
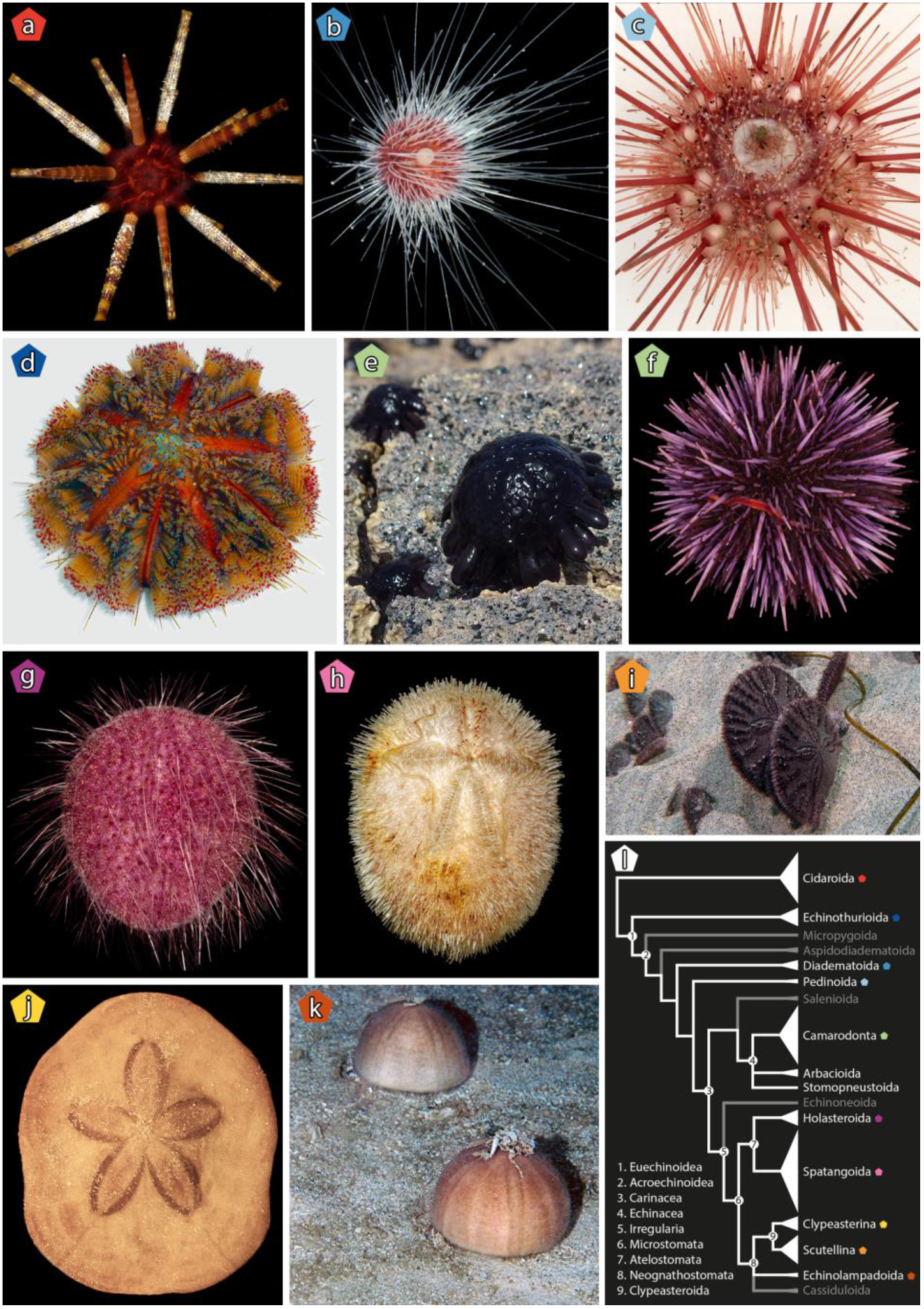
Morphological and taxonomic diversity of echinoids included in this study. **a** *Prionocidaris baculosa*. **b** *Lissodiadema lorioli*. **c** *Caenopedina havaiisensis*. **d** *Asthenosoma varium*. **e** *Colobocentrotus atratus*. **f** *Strongylocentrotus purpuratus*. **g** *Pilematechinus* sp. **h** *Brissus obesus*. **i** *Dendraster excentricus*. **j** *Clypeaster subdepressus*. **k** *Conolampas sigsbei*. **l** Current echinoid classification, modified from [6]. Clade width is proportional to the number of described extant species; clades shown in white have representatives included in this study (see Table 1). Colored pentagons are used to identify the clade to which each specimen belongs, and also correspond to the colors used in Fig. 2. Throughout, nomenclatural usage follows that of [6], in which full citations to authorities and dates for scientific names can be found. Photo credits: G.W. Rouse (**a, c, e-i**), FLMNH-IZ team (**b**), R. Mooi (**d, j**), H.A. Lessios (**k**).

The higher-level taxonomy and classification of both extant and extinct sea urchins have a long history of research (reviewed by [12, 23]). The impressive fossil record of the group, as well as the high complexity of their plated skeletons (or tests), have allowed lineages to be readily identified and their evolution tracked through geological time with a precision unlike that possible for other clades of animals (e.g., [24–28]). Morphological details of the test have also been used to build large matrices for phylogenetic analysis [9, 12, 25, 29–32]. The most comprehensive of these morphological phylogenetic analyses [12], has since served as a basis for the current taxonomy of the group (Fig. 1l). This analysis confirmed several key nodes of the echinoid tree of life that were also supported by previous efforts, such as the position of cidaroids (Fig. 1a) as sister to all other sea urchins (Fig. 1b-k)—united in the clade Euechinoidea Bronn, 1860—and the subdivision of the latter into the predominantly deep-sea echinothurioids (Fig. 1d) and the remainder of euechinoid diversity (Acroechinoidea Smith, 1981). Likewise, Kroh and Smith [12] confirmed the monophyly of some major clades such as Echinacea Claus, 1876, including all the species currently used as model organisms and their close relatives (Fig. 1e, f); and Irregularia Latreille, 1825, a group easily identified by their antero-posterior axis and superimposed bilateral symmetry [33]. The irregular echinoids were shown to be further subdivided into the extant echinoneoids, atelostomates (Fig. 1g, h) (including the heart urchins) and neognathostomates (Fig. 1i-k) (including the sand dollars). Other relationships, however, proved more difficult to resolve. For example, the pattern of relationships among the main lineages of acroechinoids received little support and was susceptible to decisions regarding character weighting, revealing a less clear cut-picture [10, 12].

In stark contrast with these detailed morphological studies, molecular efforts have lagged. Next-generation sequencing (NGS) efforts have been applied to relatively small phylogenetic questions, concerned with the resolution of the relationships within Strongylocentrotidae Gregory, 1900 [34], a clade of model organisms, as well as among their closest relatives [35]. Although several studies have attempted to use molecular data to resolve the backbone of the sea urchin phylogeny [8, 10, 25, 30, 36, 37], all these have relied on just one to three genes, usually those encoding ribosomal RNA. The lack of comprehensive sampling of loci across the genome thus limits the robustness of these phylogenies. Furthermore, recent analyses have suggested that ribosomal genes lack sufficient phylogenetic signal to resolve the deepest nodes of the echinoid tree with confidence [8].

In light of this, it is not clear how to reconcile the few—yet critical—nodes for which molecular and morphological data offer contradicting resolutions. For example, most morphological phylogenies strongly supported the monophyly of sea biscuits and sand dollars (Clypeasteroida L. Agassiz, 1835), and their origin from a paraphyletic assemblage of lineages collectively known as “cassiduloids”, including Echinolampadoida Kroh & Smith, 2010 and Cassiduloida Claus, 1880 among extant clades, as well as a suite of extinct lineages [12, 29, 31, 38]. In contrast, all molecular phylogenies to date that incorporated representatives of both groups have resolved extant “cassiduloids” nested within clypeasteroids, sister to only one of its two main subdivisions, the scutelline sand dollars [8, 10, 25, 30]. This molecular topology not only undermines our understanding of the evolutionary history of one of the most ecologically and morphologically specialized clades of sea urchins [38, 39], it also implies a strong mismatch with the fossil record, requiring ghost ranges of the order of almost 100 Ma for some clypeasteroid lineages [10, 40]. Likewise, the earliest divergences among euechinoids, including the relative positions of echinothurioids and a collection of lineages collectively known as aulodonts (micropygoids, aspidodiadematoids, diadematoids and pedinoids [41]), have consistently differed based on morphological and molecular data, often with poor support provided by both [8, 10, 12, 25, 40]. Finally, previous studies have resolved different lineages of regular echinoids, including diadematoids, aspidodiadematoids, pedinoids, salenioids and salenioids + echinaceans, as sister to Irregularia [8, 12, 25, 37].

Given the outstanding quality of their fossil record and our thorough understanding of their development, sea urchins have the potential to provide a singular basis for addressing evolutionary questions in deep-time [42], providing access to the developmental and morphological underpinnings of evolutionary innovation (e.g., [8, 43]). However, uncertainties regarding the phylogenetic history of sea urchins propagate into all of these downstream comparative analyses, seriously limiting their potential in this regard. Here, we combine available genome-scale resources with de novo sequencing of transcriptomes to perform the first phylogenomic reconstruction of the echinoid tree of life. Our efforts provide a robust evolutionary tree for this ancient clade, made possible by gathering the first NGS data for many of its distinct lineages. We then explore some important morphological transformations across the evolutionary history of the clade.

## Results

Several publicly available transcriptomic and genomic datasets are available for sea urchins and their closest relatives, the products of multiple sequencing projects [44, 45] stretching back to the sequencing of the genome of the purple sea urchin [46]. A subset of these datasets was employed here and complemented with whole transcriptomic sequencing of 17 additional species, selected to cover as much taxonomic diversity as possible. In the end, 32 species were included in the analyses, including 28 echinoids plus 4 outgroups. A complete list of these, including details on specimen sampling for all newly generated data, as well as SRA and Genome accession numbers, is provided in Table 1. All analyses were performed on a 70% occupancy matrix composed of 1,040 loci and 331,188 amino acid positions (Fig. S1), as well as constituent gene matrices.

**Table 1.** Information on the species and sequences used in the analysis. Sampling locality is shown for newly sequenced taxa, citations for data obtained from the literature. For deep-sea specimens, sampling depth is also reported.

| Clade | Species | Data type<sup>1</sup> | Sampling locality (depth)/Source | Voucher number | SRA/Genome numbers |
| --- | --- | --- | --- | --- | --- |
| Arbacioida Gregory, 1900 | *Arbacia punctulata* (Lamarck, 1816) | T | [43] | SIO-BIC E6740 | SRR2843235 |
| Camarodonta Jackson, 1912 | *Colobocentrotus atratus* (Linnaeus, 1758) | T | Kailua Kona, Hawaii Island | SIO-BIC E7012 |  |
|  | *Echinometra mathaei* (Blainville, 1825) | T | Al-Fahal Reef, Red Sea, Makkah, Saudi Arabia | SIO-BIC E6896 |  |
|  | *Evechinus chloroticus* (Valenciennes, 1846) | T | [129] | - | SRR1014619 |
|  | *Heliocidaris erythrogramma* (Valenciennes, 1846) | T | [130] | - | SRR1211283 |
|  | *Lytechinus variegatus* (Lamarck, 1816) | T | [5] | - | SRR1139214 |
|  | *Mesocentrotus nudus* (A. Agassiz, 1864) | T | [131] | - | SRR5017175 |
|  | *Paracentrotus lividus* (Lamarck, 1816) | T | [132] | - | SRR1735501 |
|  | *Sphaerechinus granularis* (Lamarck, 1816) | T | [5] | - | SRR1139199 |
|  | *Strongylocentrotus purpuratus* (Stimpson, 1857) | G | [45]; Spur_4.2 | - | GCF 000002235.4 |
| Cidaroida Claus, 1880 | *Eucidaris tribuloides* (Lamarck, 1816) | T | [43] | SIO-BIC E6742 | SRR2844625 |
|  | *Prionocidaris baculosa* (Lamarck, 1816) | T | Al-Fahal Reef, Red Sea, Makkah, Saudi Arabia | SIO-BIC E6897 |  |
| Clypeasteroida sensu A. Agassiz, 1872 | *Clypeaster rosaceus* (Linnaeus, 1758) | T | Bocas del Toro, Panama | - |  |
|  | *Clypeaster subdepressus* (Gray, 1825) | T | Bocas del Toro, Panama | - |  |
|  | *Dendraster excentricus* (Eschscholtz, 1831) | T | [43] | SIO-BIC E5640 | SRR2844623 |
|  | *Echinarachnius parma*<br>(Lamarck, 1816) | T | [5] | - | SRR1139193 |
|  | *Echinocyamus crispus* Mazetti,<br>1893 | T | Al-Fahal Reef, Red Sea, Makkah,<br>Saudi Arabia | SIO-BIC E6903 |  |
|  | *Mellita tenuis* H.L. Clark, 1940 | T | Apalachee Bay, Wakulla County,<br>Florida | SIO-BIC E7015 |  |
| Diadematoida<br>Duncan, 1889 | *Diadema setosum* (Leske, 1778) | T | Al-Fahal Reef, Red Sea, Makkah,<br>Saudi Arabia | SIO-BIC E6905 |  |
|  | *Lissodiadema lorioli* Mortensen,<br>1903 | T | Kaneohe Bay, Honolulu, Hawaii<br>Island | UF Echino<br>18893 |  |
| Echinolampadoida<br>Kroh & Smith, 2010 | *Conolampas sigsbei* (A.<br>Agassiz, 1878) | T | Willemstad, Curaçao (233-300 m) | - |  |
| Echinothurioida<br>Claus, 1880 | *Araeosoma leptaleum* A.<br>Agassiz & H.L. Clark, 1909 | T | Mount Quepos, Pacific Ocean,<br>Costa Rica (1097 m) | SIO-BIC E7021 |  |
|  | *Asthenosoma varium* Grube,<br>1868 | T | Momi Bay, Viti Levu, Fiji | - |  |
| Holasteroida Durham<br>& Melville, 1957 | *Pilematechinus* sp. | T | Axial Seamount, Juan de Fuca<br>Ridge (1550 m) | SIO-BIC E6947 |  |
| Pedinoida<br>Mortensen, 1939 | *Caenopedina hawaiiensis* H.L.<br>Clark, 1912 | T | Mount Quepos, Pacific Ocean,<br>Costa Rica (1908 m) | SIO-BIC E7020 |  |
| Spatangoida L.<br>Agassiz, 1840 | *Brissus obesus* Verrill, 1867 | T | San Clemente Island, California | SIO-BIC E7018 |  |
|  | *Meoma ventricosa* (Lamarck,<br>1816) | T | Bocas del Toro, Panama | - |  |
| Stomopneustoida<br>Kroh & Smith, 2010 | *Stomopneustes variolaris*<br>(Lamarck, 1816) | T | Sohoa, Mayotte | SIO-BIC E7014 |  |
| Holothuroidea de<br>Blainville, 1834 | *Holothuria forskali* Delle<br>Chiaje, 1823 | T | [133] | - | SRR5109955 |
| Asteroidea de<br>Blainville, 1830 | *Acanthaster planci* (Linnaeus,<br>1758) | G | [134]; OKI-Apl_1.0 | - | GCA<br>001949145.1 |
|  | *Patiria miniata* (Brandt, 1835) | G | [44]; Pmin_1.0 | - | GCA<br>000285935.1 |
| Hemichordata<br>Bateson, 1885 | *Saccoglossus kowalevskii* A.<br>Agassiz 1873 | G | [135]; Skow_1.1 | - | GCA<br>000003605.1 |

<sup>1</sup> T = transcriptome, G = genome.

Initial analyses were complicated by the problem of resolving the position of *Arbacia punctulata;* different methods resolved this species as either a member of Echinacea, as suggested by previous morphological and molecular studies [10, 12], or as the sister lineage to all remaining euechinoids. Further analyses suggested that this second, highly conflicting topology might be the consequence of sequence contamination (see Fig. S2). Topologies obtained after attempting to control this problem showed strong support for a monophyletic Echinacea—including *Arbacia puntulata, Stomopneustes variolaris* and camarodonts—although the relationships among these three lineages received only weak support (Fig. S2). Given the ad hoc nature of our approach, we regard this result as preliminary, and excluded *Arbacia* from all subsequent analyses.

Phylogenomic matrices are the product of complex evolutionary histories which are only partially captured by our current models of molecular evolution. This often results in fully supported yet incorrect topologies, as all methods are susceptible to systematic biases in various ways and to different degrees [47, 48]. In order to explore the effects of model selection, phylogenetic inference was performed on the concatenated alignment using a diversity of procedures, including maximum likelihood (ML) inference using two different mixture models and the best-fit partitioning scheme, as well as Bayesian inference (BI) under site-homogenous and site-heterogenous models (see Methods). All five methods of probabilistic inference recovered exactly the same phylogeny (Fig. 2a), showing the robustness of our results to the implementation of different approaches to model molecular evolution. Furthermore, support was maximum for almost all nodes across all methods, and no other tree was found in the credible set of topologies explored by either of the BI methods. This phylogeny shows strong agreement with the current higher-level classification of echinoids, supporting the monophyly of most previously recognized clades classified at or above the level of order. These include the position of Cidaroida as sister to all other echinoids, the monophyly and close relationship of Echinacea and Microstomata (including all sampled irregular echinoids), and the subdivision of the latter into atelostomates and neognathostomates (as labelled on the tree, Fig. 2a). Relationships at lower taxonomic levels are beyond the scope of this study, as only one or two species per major clade were sampled, with the exception of camarodonts and scutelline sand dollars. Internal relationships among camarodonts fully agree with recently published estimates based on mitochondrial genomes [35], even though our taxonomic sampling differs. For scutelline sand dollars, our phylogeny confirms a close relationship of Dendrasteridae to Echinarachniidae, as suggested by early DNA hybridization assays [49], rather than between Dendrasteridae and Mellitidae, as previously argued based on morphological evidence [9, 12, 38].

**Fig. 2.**
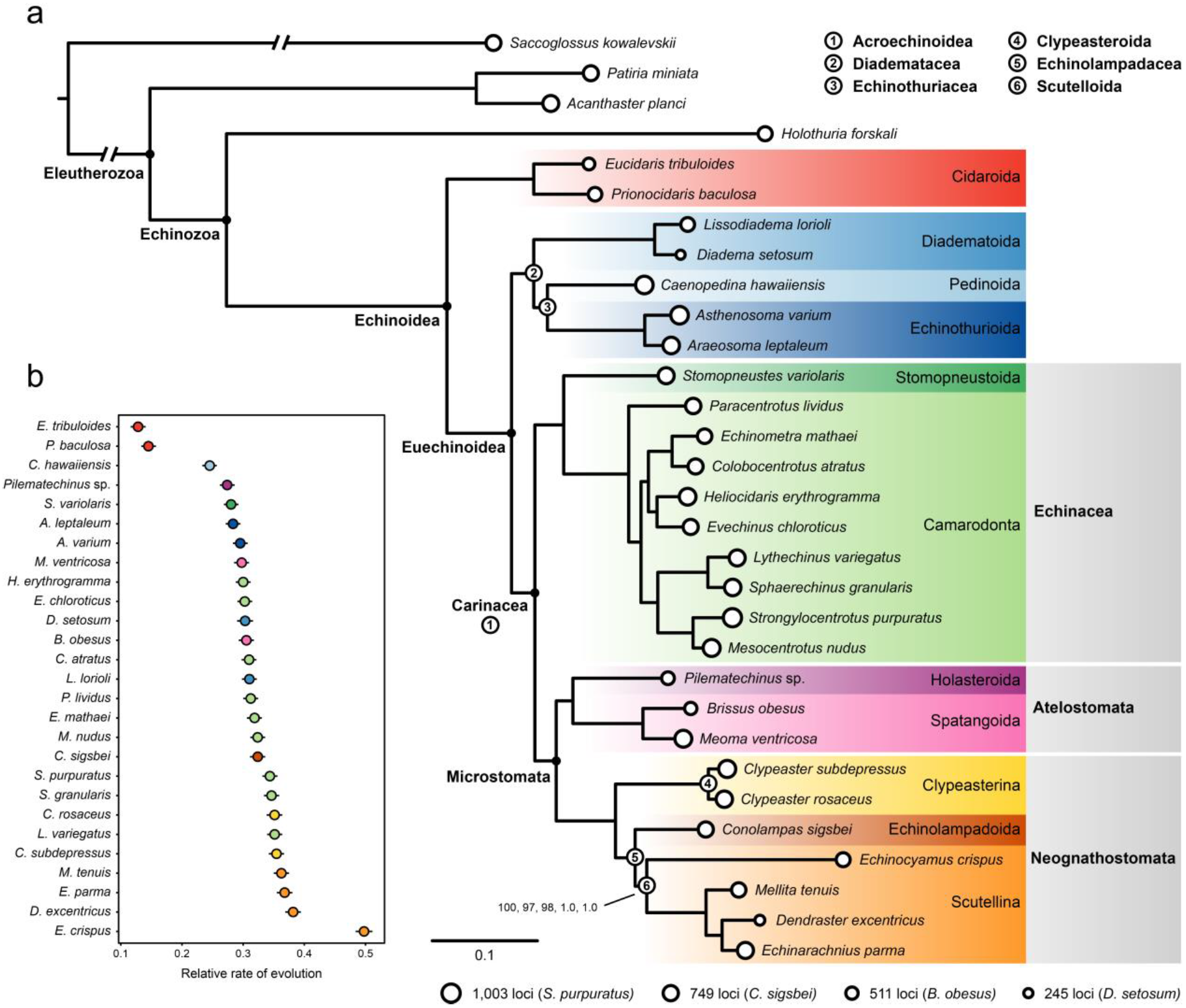
**a** Maximum likelihood phylogram corresponding to the unpartitioned analysis. The topology was identical across all four probabilistic methods employed, and all nodes attained maximum support except for the node at the base of Scutellina, which received a bootstrap frequency of 97 and 98 in the maximum likelihood analyses under the LG4X and PMSF mixture models, respectively (see Methods). Circles represent number of genes per terminal. Numbered nodes denote novel taxon names proposed or nomenclatural amendments (see Discussion), and are defined on the top right corner. **b** Distance of each ingroup species to the most recent common ancestor of echinoids, which provides a metric for the relative rate of molecular evolution. Dots correspond to mean values out of 2,000 estimates obtained by randomly sampling topologies from the post burn-in trees from PhyloBayes (using the CAT-Poisson model), which better accommodates scenarios of rate variation across lineages. Lines show the 95% confidence interval.

On the other hand, our topology conflicts with current echinoid classification (Fig. 1l) in two main aspects. First, it does not recover echinothurioids as sister to the remaining euechinoids, therefore contradicting the monophyly of Acroechinoidea. Instead, Echinothurioida is recovered as a member of a clade that also incorporates the lineages of aulodonts that were sampled—pedinoids and diadematoids (Fig. 1b, c). Second, and more surprisingly, it rejects the monophyly of the sea biscuits and sand dollars, proposing instead a sister relationship between *Conolampas sigsbei* (an echinolampadoid) and only one of the two main subdivisions of clypeasteroids, the scutellines. Both of these topologies were recovered by previous molecular analyses [8, 10, 25], but were disregarded due to the perceived strong conflict with morphological data [12, 40].

We further explored coalescent-based inference using ASTRAL-II [50], which recovered a very similar topology to the other approaches. Notably, however, it strongly supported the placement of *Conolampas* in an even more nested position, inside the clade formed by scutelline sand dollars, sister to Scutelliformes Haeckel, 1896 (Fig. 3a). Exploration of gene tree incongruence using a supernetwork approach revealed topological conflicts among gene trees in the resolution of the *Conolampas* + scutelline clade, with *Conolampas, Echinocyamus* and scutelliforms forming a reticulation (Fig. 3a, inset). We hypothesize this to be the consequence of high levels of gene tree error caused by the heterogeneity in rates of evolution among the included lineages, with *Conolampas* evolving significantly slower, and *Echinocyamus* significantly faster, than the scutelliforms (as shown by non-overlapping 95% confidence intervals in Fig. 2b). To test this hypothesis, we performed species tree inference with ASTRAL-II using approximately a third of the gene trees, selecting those derived from genes with the lowest levels of both saturation and rate heterogeneity across lineages (Fig. 3c). The resulting topology agrees with those from the other methods in every detail, with the position of *Conolampas* shifting to become sister to the scutellines with a relatively strong local posterior probability (localPP) of 0.91 (Fig. 3b). In contrast, most species trees derived from equal-sized subsets of randomly selected gene trees provide strong support for placing *Conolampas* in disagreement with the position obtained by other methods (average localPP = 0.92; Fig. 3d). The few replicates in which *Conolampas* is recovered as sister to the scutellines (16%), receive low support values (average localPP = 0.51; Fig. 3d).

**Fig. 3.**
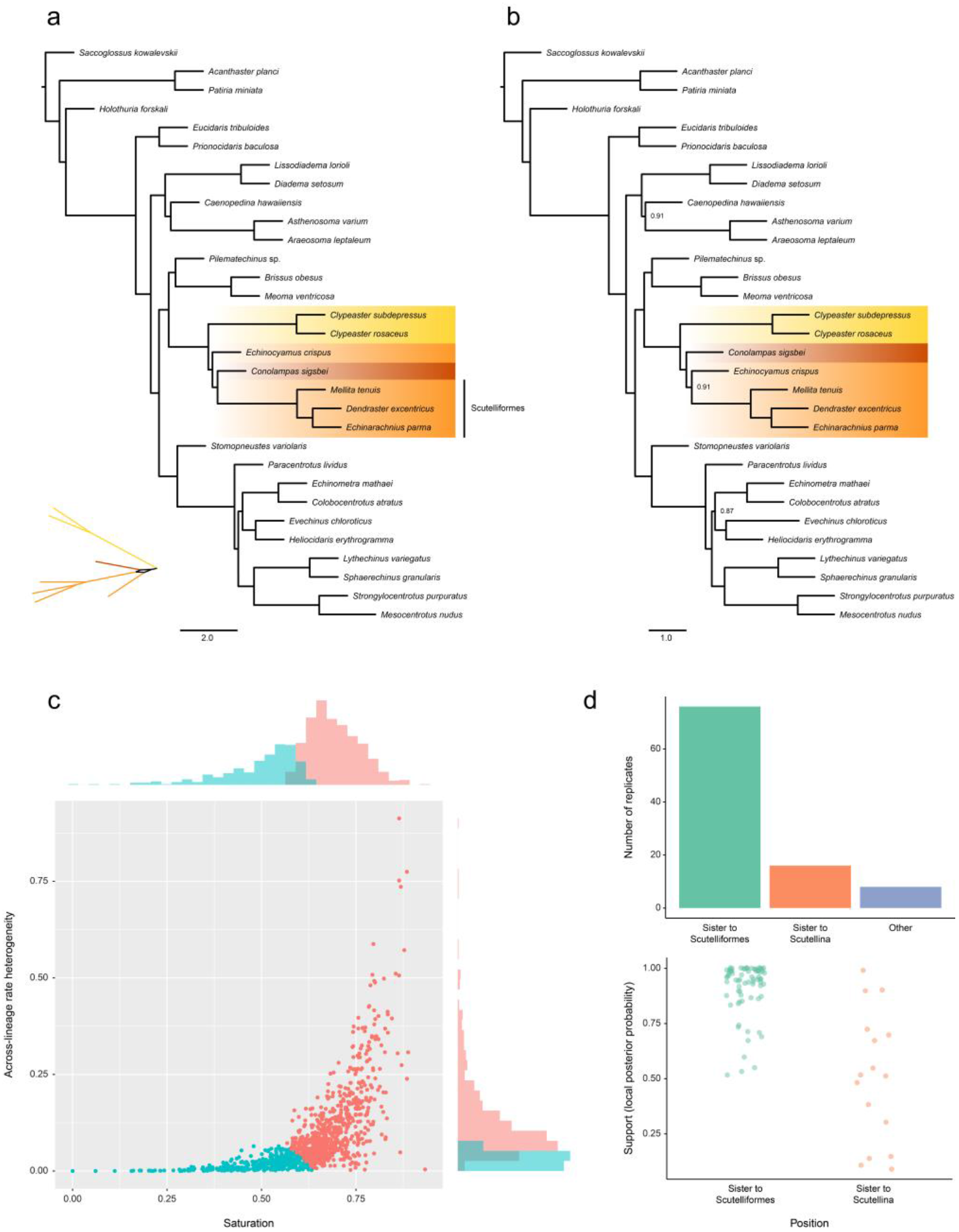
Phylogenetic inference using the coalescent-based summary method ASTRAL-II. **a** Phylogeny obtained using all 1,040 gene trees. The phylogeny conflicts with that obtained using all other methods by placing *Conolampas sigsbei* inside Scutellina, sister to Scutelliformes. The neognathostomate section of a supernetwork built from gene tree quartets is also depicted, showing a reticulation involving *Conolampas, Echinocyamus* and scutelliforms. **b** Phylogeny obtained using 354 gene trees, selected to minimize the negative effects of saturation and across-lineage rate heterogeneity. The position of *Conolampas* shifts to become sister to Scutellina (as in all other methods), with relatively strong support. To emphasize the shift in topology between the two, only neognathostomate clades have been colored (as in Fig. 2), and nodes have maximum local posterior probability unless shown. **c** Values of the two potentially confounding factors across all genes. Genes in red were excluded from the analysis leading to the topology shown in b. Histograms for both variables are shown next to the axes. **d** Summary of the results obtained performing inference with ASTRAL-II after deleting 66% of genes selected at random (100 replicates). Most replicates showed the same topology as in **a**. Only 16% placed *Conolampas* as sister to Scutellina (top), and even among them the support for this resolution was generally weak (bottom).

Finally, we used a series of topological tests to assess the strength of evidence for our most likely topology against the two traditional hypotheses of relationships with which it conflicts most strongly: the monophyly of Acroechinoidea and Clypeasteroida, clades that are supported by morphological data [12] and recognized in the current classification of echinoids (Fig. 1l). SOWH tests [51] strongly rejected monophyly in both cases (both *P* values < 0.01). We were able to trace the signal opposing the monophyly of these two clades down to the gene level, with a predominant fraction of genes showing support for the novel position of Echinothurioida united with Pedinoida and Diadematoida, as well as for the position of echinolampadoids as sister to the scutelline sand dollars (Fig. 4). Genes supporting these novel groupings showed strong preference for them, while the comparatively smaller fraction of genes favoring the traditional resolutions did so only weakly

**Fig. 4.**
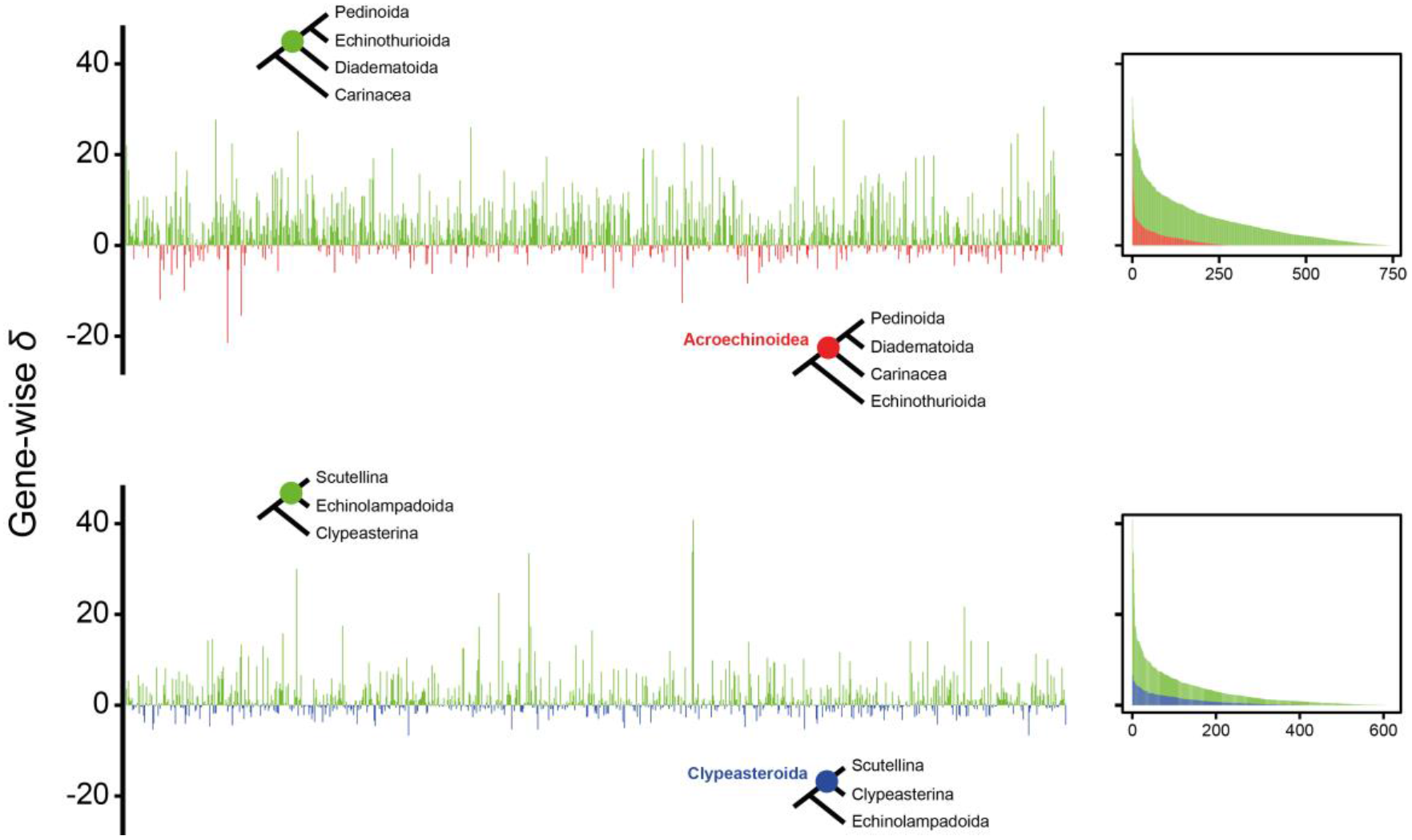
Distribution of phylogenetic signal for novel resolutions obtained in our phylogenomic analyses. Signal is measured as the difference in gene-wise log-likelihood scores (*δ* values) for the unconstrained (green) and constrained topologies enforcing monophyly of Acroechinoidea (top, red) or Clypeasteroida (bottom, blue). The same results are shown on the right, except that values are expressed as absolute differences and genes are ordered following decreasing *δ* values to show the overall difference in support for both alternatives.

Furthermore, we were unable to detect any evidence that this signal arises from non-historical sources. Gene-wise *δ* values (i.e., the difference in log-likelihood score for constrained and unconstrained ML topologies for each individual locus) showed no correlation with several potentially biasing factors, including compositional heterogeneity, among-lineage rate variation, saturation, and amount of missing data (multiple linear regression, *P* = 0.167 and 0.165 for *δ* values obtained constraining clypeasteroid and acroechinoid monophyly, respectively; see Fig. 5). Thus, we detect no evidence that the support for these novel hypotheses stems from anything other than phylogenetic history.

**Fig. 5.**
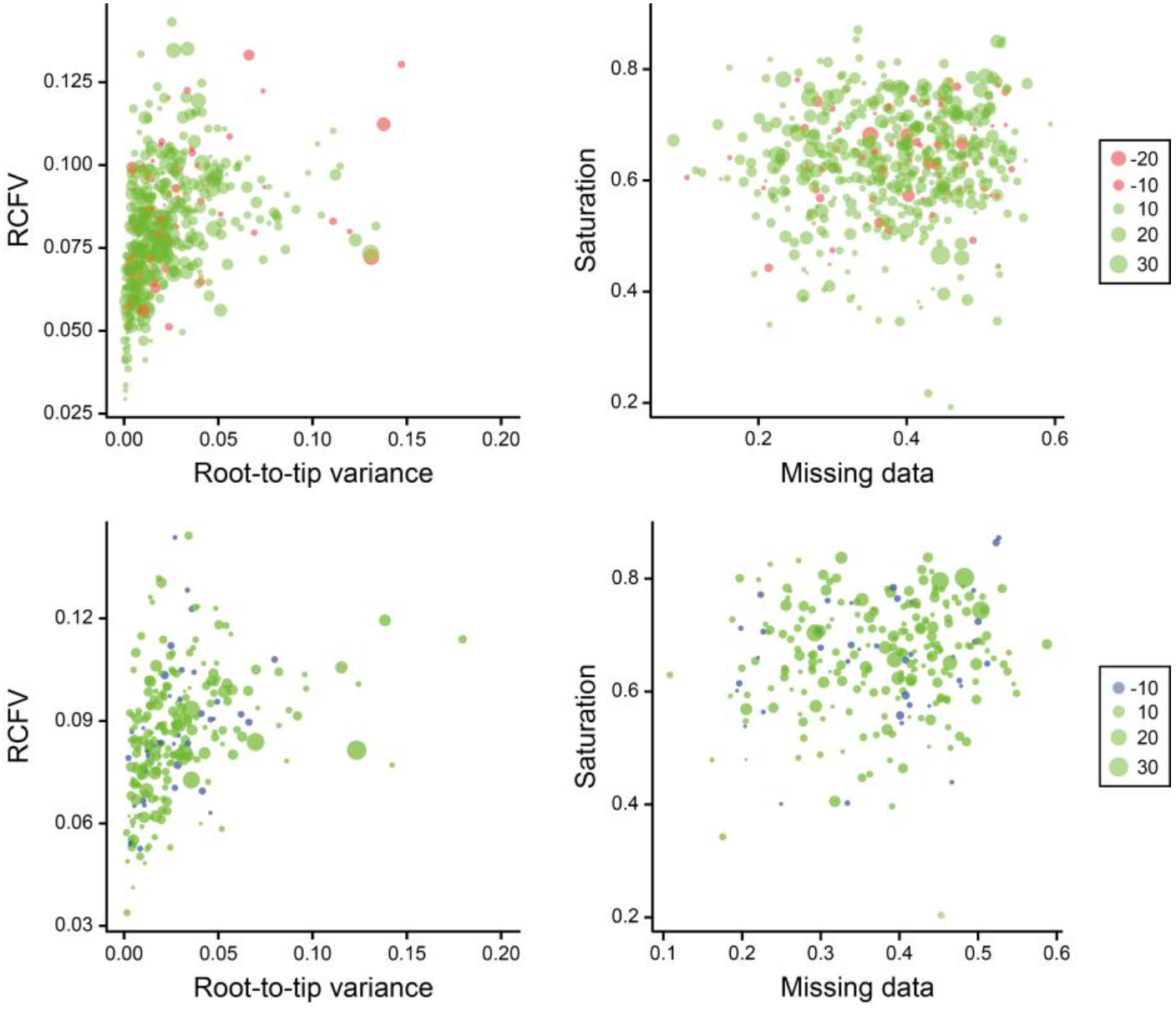
Exploration of potential non-phylogenetic signals biasing inference. Gene-wise *δ* values (as shown in Fig. 4) obtained by constraining acroechinoid (top) and clypeasteroid (bottom) monophyly are shown using dot size and color (see legend). Root-to-tip variance axis was truncated to show the region in which most data points lie. The relative support for these topological alternatives is not correlated with any of the four potentially biasing factors explored, as seen by the lack of clustering of genes with similar *δ* values along the axes.

## Discussion

### General comments

Since the publication of Mortensen’s seminal monographs (starting almost a century ago), echinoid classifications have largely relied on morphological data. Detailed study of the plate arrangements in the echinoid test has proved a rich source of characters for both fossil and extant taxa, integrating them in a unified classification scheme. However, the amount of time separating the main echinoid lineages, coupled with the profound morphological reorganization they have experienced, have resulted in parts of their higher-level classification remaining uncertain. Although molecular data offer an alternative source of phylogenetic information, efforts so far have largely targeted a restricted character set of limited utility for deep-time inference, resulting in issues similar to those faced by morphological attempts. Phylogenomics hold the potential to provide insights into the deep evolutionary history of echinoids, an avenue explored here for the first time.

Analysis of our phylogenomic dataset provided similar estimates of phylogeny using either concatenation or coalescent-based methods (Figs. 2a and 3a), with the exception of one node that was resolved differently by the two approaches. This node involved the order of divergences among two lineages with dissimilar rates of molecular evolution, namely *Echinocyamus crispus*, a scutelline sand dollar with the fastest rate of evolution among all sampled taxa, and *Conolampas sigsbei*, a relatively slow-evolving echinolampadoid (at least in the context of the remaining neognathostomates, Fig. 2b). Extensive rate variation among neognathostomate lineages has been reported previously, with potential consequences for phylogenetic inference and time-calibration [25, 27, 48]. Although increased taxonomic sampling is required, several lines of evidence suggest that the tree obtained by ASTRAL-II is artefactual, including the strong support for the alternative resolution found by all other methods, the implausible morphological history that this species tree implies, and its even greater departure from previous phylogenetic results [10, 25, 30]. We were able to bring ASTRAL-II into agreement with concatenation-based approaches by including only those genes expected to better handle the difference in evolutionary rate among the sampled taxa (Fig. 3b, c).

The resulting topology shows strong support for the same resolution of Neognathostomata found across all concatenation-based approaches. Although we did not formally test the reason behind this change in topology inferred with ASTRAL-II, we found that species trees obtained from randomly subsampled gene trees are generally identical to the one supported by the full set of gene trees (Fig. 3d). The widespread adoption of methods accounting for incomplete lineage sorting is one of the major innovations made possible by phylogenomics [52], but its utility for phylogenetic inference in deep time remains a topic of discussion [53, 54]. Simulations have demonstrated that genes with minimal phylogenetic information might produce unreliable gene trees, which in turn reduce the accuracy of gene tree estimation using summary methods [55, 56]. Our empirical analysis shows that rate heterogeneity among neognathostomate echinoids might be strong enough to bias some coalescent-based approaches, potentially by reducing the phylogenetic signal of individual genes and affecting gene tree accuracy.

The topology of the tree obtained using probabilistic methods of inference (Fig. 2a) is consistent in many ways with previous analyses of Echinoidea, both morphological and molecular. The two cidaroids sampled are recovered as sisters, and as a clade they are joined to all other echinoids at the earliest internal node of the group; Euechinoidea is thus supported by our findings. Some previous authors relied heavily on the fact that adults of some irregular echinoid taxa have an Aristotle’s lantern (e.g., clypeasteroids), while others lack the jaw apparatus entirely (e.g., spatangoids and holasteroids), to argue that Irregularia is polyphyletic [57–59]. These arguments have since been rejected by nearly every phylogenetic analysis [9, 10, 12, 25, 29–31, 60–62]; the strong support in our phylogenomic analysis for the monophyly of the sampled irregular echinoids is therefore largely uncontroversial. Although our taxonomic sampling is insufficient to establish which clade constitutes the sister group to Irregularia, we do not recover diadematoids or pedinoids in such a position, as previously suggested [12, 37]. Instead, our topology shows Echinacea as their closest relative among the sampled taxa (as in [25], among others). Within irregular echinoids, Atelostomata von Zittel, 1879 has long been regarded as monophyletic, comprising two major extant clades, holasteroids and spatangoids [12, 63], a topology further supported by previous molecular analyses (e.g., [25]). Taxon sampling for the deep-sea holasteroids continues to be a challenge, but we were able to sample what we have determined to be a new species of *Pilematechinus*. Our phylogenomic analysis strongly supports a sister group relationship between this holasteroid and two species of brissid spatangoids, which themselves form a clade.

There remain two major points of departure between our phylogenomic tree (regardless of method choice) and those generally accepted. One discrepancy concerns the echinothurioid and aulodont taxa. The other involves the “cassiduloids” and clypeasteroids (sensu lato). We find maximum support for novel resolutions of these clades among all probabilistic methods explored, including both site-homogenous and heterogenous approaches to model molecular evolution. These rely on different underlying assumptions and are able to cope with problems such as saturation and rate variation to different extents, thus often producing contradicting topologies [47, 64, 65]. SOWH topological tests show that our phylogenomic data significantly reject the traditional resolution of these clades. We find no evidence that this signal is restricted to a few “outlier” genes or that it stems from systematic biases (as is the case with many phylogenomic datasets, e.g., [66–70]), but rather appears to be the result of true phylogenetic signal distributed throughout the genome (Figs. 4 and 5).

Should further testing with an expanded taxonomic sampling support the topology of our tree it will have significant implications for echinoid research, systematics, and paleontology. We examine these implications below to explore how they can be reconciled with former and present views of the evolution of the groups in question and to propose appropriate nomenclatural changes.

### Non-monophyly of Acroechinoidea sensu Smith, 1981

Our result departs from that of nearly every recent morphological analysis in placing both of our sampled, distantly related diadematoids (sensu [12]) as sister to a clade uniting echinothurioids and pedinoids. Notably, previous molecular studies had found a clade composed of these three lineages (e.g., [25]), a result that was not explored because this clade was not recovered in a total evidence inference incorporating morphological data.

There are several morphological similarities that could be interpreted as evidence of a relationship between echinothurioids and diadematoids [12, 29, 71]. In spite of these, a purported lack of “advanced” features was deemed to make echinothurioids too unlike other euechinoids, ultimately leading to their placement as sister to the remainder of euechinoid diversity (Acroechinoidea). This topology was counter to earlier classifications, notably that of Durham and Melville [57], who placed pedinids with echinothuriids (both at the family level) in the order Echinothurioida, which they united with Diadematoida into Diadematacea Duncan, 1889. This is precisely the hierarchical arrangement recovered by our phylogenomic analysis.

In proposing Acroechinoidea, Smith [9] listed several plesiomorphies of echinothurioids that made them the “primitive sister group to all other euechinoids” (p. 792) including the imbricate, flexible test, ambulacral plate columns that extend onto the peristomial membrane (i.e., lack of specialized buccal plates [72]), internal coelomic pouches associated with the lantern (Stewart’s organs), a somewhat flattened lantern with a U-shaped foramen magnum in the pyramids, and shallow, grooved teeth. Kroh and Smith [12] specifically mentioned two features of acroechinoids supporting their monophyly: plate compounding, with ambulacral primary tubercles mounted on more than one plate; and reduction of the ambulacral plating on the peristomial membrane to 5 pairs of buccal plates.

Flexibility of the test corona is ubiquitous among Triassic forms such as miocidarids, thought to have given rise to all post-Paleozoic echinoids [73]. Test rigidity would have had to evolve independently in cidaroids and acroechinoids for the echinothurioid condition to represent a retention of this plesiomorphy. We suggest instead that flexibility originated secondarily among echinothurioids, possibly as an adaptation to the difficulties of secreting calcium carbonate in the deep-sea. Echinothurioid plate morphology, arrangement, and distribution of collagen between the plates are unlike those in Paleozoic forms, supporting this interpretation. Furthermore, imbrication and slight flexibility are also present in coronal regions of some diadematoids.

Stewart’s organs are present in cidaroids, echinothurioids and some diadematoids, with vestigial remnants in pedinoids [9]; the loss of this organ cannot therefore constitute an acroechinoid synapomorphy. Likewise, although the ambulacral plating in echinothurioids is unusual among euechinoids, it is fully consistent with the diadematoid pattern of triplets [74] and cannot be the basis for removing echinothurioids from the acroechinoids (sensu [9]). Even though echinothurioids differ from diadematoids by lacking a primary tubercle spanning the triplets, this could be related to the overall spine size reduction among echinothurioids.

Monophyly of acroechinoids has been supported previously by citing loss of ambulacral plating on the peristomial membrane, where only five ambulacral pairs of buccal plates are present [9, 12]. However, the aberrant *Kamptosoma* (see [71, 74]), recently suggested to be sister to all other echinothurioids [12], is unique among them in having five pairs of buccal plates, just as in acroechinoids. *Kamptosoma* retains an apparently plesiomorphic condition not only for this clade, but for all euechinoids, whereas all other echinothurioids possess an autapomorphic, plated peristomial membrane. Fully consistent with this, the peristomial regions of cidaroids and echinothurioids grow in substantially different ways [75, 76], suggesting that the continuation of ambulacra onto the peristomial membrane is not homologous between them. *Kamptosoma* also has a similar structure of tooth plates to that of diadematoids [71], and possesses crenulate tubercles, otherwise absent from echinothurioids [74]. The spines of diadematoids and echinothurioids tend to be either hollow or have a lumen filled with reticulated stereom [77]. Although most pedinoids have solid primary spines, those in *Caenopedina* also have reticulated stereom in the lumen (Coppard SE, unpublished data). Diadematoids and echinothurioids are also notorious as the only echinoids with venom-bearing spines, a potential synapomorphy that unites them in the same clade.

Diadematoids, echinothurioids and pedinoids likely diverged from each other sometime during the Triassic [8, 40], and the length of ensuing time has obscured their commonalities, as already noted by Mortensen [78]. Further analysis of the ontogeny of echinothurioids is needed to determine how their unique features are gained, or how apomorphies attributed to acroechinoids (sensu lato) might have been lost. However, almost no ontogenetic information exists for echinothurioids or pedinoids. Chemical analysis of the venoms in echinothurioids and diadematoids could be used to test whether these systems are homologous, exploring whether less robust test development is related to enhanced protection afforded by venomous spination. The relationship of these features to abyssal environments is also poorly understood. Future phylogenomic work is needed to place the remaining aulodont lineages (aspidodiadematoids and micropygoids) within this novel phylogenetic structure.

The topology of our tree implies nomenclatural changes to the current echinoid classification scheme [12]. Extending the concept of acroechinoids to include echinothurioids would be redundant with the concept of Euechinoidea, and counter to the original concept of Acroechinoidea presented by Smith [9]. Restricting Acroechinoidea to all euechinoids except for echinothurioids, diadematioids, and pedinoids would make the junior term Carinacea redundant. In accepting the topology presented herein, we abandon the term Carinacea, and amend Acroechinoidea to include all euechinoids other than diadematoids + echinothurioids + pedinoids. For this latter group, we resurrect the name Diadematacea Duncan, 1889 (sensu [57]). If a name is needed for the pedinoid + echinothurioid clade, we recommend using Echinothuriacea.

### Non-monophyly of Clypeasteroida L. Agassiz, 1835

Sand dollars and sea biscuits (clypeasteroids) are recognizable at a glance and their monophyly has been resoundingly supported by all morphological analyses (e.g., [9, 12, 38, 60, 79–81]). According to Kroh and Smith [12], the clade contains two subgroups: Clypeasterina L. Agassiz, 1835 (sea biscuits), and Scutellina Haeckel, 1896, including scutelliforms (“true” sand dollars) and laganiforms (sea peas and sun dollars). Clypeasteroids (sensu lato) are presently grouped with so-called “cassiduloids” in the clade Neognathostomata Smith, 1981. Among these, the oligopygids have been considered sister to clypeasteroids [12, 38, 39, 82, 83], as both share the presence of a lantern as adults, a trait otherwise absent among neognathostomates. However, lanterns are present in juvenile forms of members of all the main extant “cassiduloid” clades [76, 84, 85], and their teeth are very similar to those of clypeasterines and scutellines [85]. Such similarities suggest that lanterns in adult clypeasterines, scutellines and oligopygoids represent the re-expression in later ontogenetic stages of a trait never fully lost [38, 60].

Littlewood and Smith [30] supported a monophyletic Clypeasteroida based on a total evidence approach, even though their rRNA tree showed a cassidulid within clypeasteroids in a sister group relationship to Scutellina. This result was also obtained by Smith et al. [25] in an analysis including multiple “cassiduloids”, all of which grouped together as sister to the scutellines, with clypeasterines again falling sister to this clade. Smith [40] indicated that there was no evidence to suggest this topology was the result of biases but conceded that “it is hard to reconcile this observation with the strong morphological evidence for clypeasteroid monophyly” (p. 304).

Nor can we attempt a full reconciliation here. However, we can suggest ways of explaining these results in light of the strong support for clypeasteroid non-monophyly in our phylogenomic analysis. Mooi [38] noted that the lantern supports of scutellines and clypeasterines are dramatically different, with the configuration present in scutellines being entirely unique to that group. This could be reinterpreted as an indication that lanterns reappeared in adults separately in the two clades. The presence of lanterns in juvenile “cassiduloids” implies that the genetic architecture associated with the lantern was never lost from the ancestors of either the clypeasterines or scutellines, allowing this transition to occur multiple times. There are other significant differences between the lanterns of clypeasterines and scutellines (illustrated in [86]) that could also be explained by a separate derivation of the structure in the two lineages. We therefore suggest this trait might not constitute a clypeasteroid synapomorphy.

Mooi [38] listed several other features supporting monophyly of clypeasteroids, but in almost every case, there are substantial differences in the way the features are expressed in clypeasterines and scutellines. For example, the number of sphaeridia is reduced in both, yet their morphology and degree of reduction is entirely different. Although these differences were originally interpreted as part of a transition series, they could also be evidence of non-homology. In terms of their ecology, clypeasterines exploit more specific food sources than scutellines [87–89], such as Foraminifera and other dominant species of infauna and epifauna (Mooi R, unpublished data), again implying that the similarities between clypeasterines and scutellines might be superficial, driven by commonalities in their modes of life.

However, there are two major features that are shared by clypeasterines and scutellines, absent not just in “cassiduloids”, but throughout most of the remainder of Echinoidea. One is the set of internal buttresses and pillars inside the test, and the other is the enormous multiplication of tube feet throughout the ambulacra. Both of these features were cited by Seilacher [90] as part of the “sand dollar paradigm”— adaptations of greatly flattened echinoids to reduced exposure to drag forces and increasing the efficiency of podial particle picking [87]. It is possible that these major, shared features of clypeasterines and scutellines are also convergences driven by adaptation to life on shifting sediments in hydrodynamically active environments. There are many other examples of parallel evolution among echinoids in response to similar evolutionary challenges [91], such as the postulated independent origin of several “sand dollar features” in arachnoidids and scutelliforms [38, 90], and the presence of internal buttresses in discoidid holectypoids [12, 92]. These morphologies constitute important avenues for further analysis in view of the non-monophyly of clypeasteroids.

Although it remains possible that sand dollar features were lost in “cassiduloids”, such a reversal is likely even less parsimonious, implying more evolutionary events than the convergent appearance of these features in clypeasterines and scutellines, especially when fossil taxa are considered. A more robust resolution of the relationships of clypeasterines and scutellines to oligopygids and other “cassiduloids” is needed to constrain these evolutionary scenarios. If the results from all recent molecular work can be taken at face value, not just the echinolampadoids, but cassiduloids, and possibly even apatopygids [39] are part of the sister clade to the scutellines. Consequently, both clypeasterines and scutellines may have originated much earlier than once hypothesized, at least prior to the Cenozoic, and possibly even in the early Cretaceous. In light of our phylogenomic topology, a reinterpretation of the morphology of the lantern system (including the arrangement of the lantern supports) might suggest that the extinct oligopygids are sister to clypeasterines alone. Further work on the morphology of the lantern present in early developmental stages of extant “cassiduloids” should provide a test of the independent derivation of lantern types and establish the likelihood of various scenarios implied by clypeasteroid non-monophyly.

Clypeasteroida derives its name from Clypeasterina. Therefore, the clade that now contains scutelliforms + laganiforms requires a new name, and we propose Scutelloida. As yet, we do not know all the successive outgroups to this clade, and whether it includes all, or a subset of the “cassiduloids”. However, at minimum from the topology of the phylogenomic tree, we recognize a clade that includes Echinolampadoida + Scutelloida, named Echinolampadacea.

## Conclusions

This study expands the set of transcriptomic resources for sea urchins, providing the first available data for many distinct lineages. Phylogenetic analyses of the resulting datasets provide a robust resolution of the backbone of the echinoid tree of life, settling many uncertainties regarding the position of multiple clades. Our efforts resolve several conflicting nodes among previous morphological and molecular approaches in favor of the latter and represent a major step towards unravelling the evolutionary history of this ancient clade. Further work is required to confirm the placement of some remaining lineages within this novel topology, as well as to interpret fully its evolutionary implications, especially with respect to the implied morphological convergences between sand dollars and sea biscuits. Nonetheless, our phylogenomic study opens up new lines of research exploring the evolution of morphology and development among sea urchins in a phylogenetically explicit framework.

## Methods

Publicly available datasets were downloaded from either NCBI or EchinoBase [45]. Although substantial genomic resources have been recently gathered for echinoids, most of these come from relatively closely related species, and sampling of the main echinoid lineages remains sparse. From the available data, we chose to include four high-quality genomes and 11 transcriptomic datasets (Table 1). All transcriptomes were obtained using pair-end sequencing in Illumina platforms, with a sequencing depth of no less than 18 million reads (average = 29.7 million). These were supplemented with 17 newly sequenced species, significantly increasing coverage of the main lineages of echinoids. Total RNA was extracted from either fresh tissues or from tissues preserved in RNA*later* (Invitrogen) buffer solution. For large specimens, tissue sampling was restricted to tube feet, muscles and/or gonads, so as to avoid contamination with gut content. Whenever possible, small specimens were starved at least overnight, before extraction of total RNA from whole animals. Extractions were performed using Ambion PureLink RNA Miniprep Kit (Life Technologies) or Direct-zol RNA Miniprep Kit (with in-column DNase treatment; Zymo Research) from Trizol. mRNA was isolated with Dynabeads mRNA Direct Micro Kit (Invitrogen). RNA concentration was estimated using Qubit RNA broad range assay kit (average 76.1 ng/μL, range = 36.6 − 166), and quality was assessed using RNA ScreenTape with an Agilent 4200 TapeStation or total RNA Nano Chips on an Agilent Bioanalyzer 2100. Values were used to customize downstream protocols following manufacturers’ instructions. Library preparation was performed with either Illumina TruSeq RNA or KAPA-Stranded RNA-Seq kits, targeting an insert size in the range of 200–300 base pairs (bp). Quality, concentration and molecular weight distribution of libraries were assessed using a combination of DNA ScreenTape, a Bioanalyzer 2100 and KAPA (qPCR-based) library quantification kits. Libraries were sequenced in multiplexed pair-end runs using Illumina HiSeq 2500 or 4000 platforms, with between 2 and 8 libraries per lane, resulting in an average sequencing depth of 49.5 million reads (range: 38.0 – 88.6). In order to minimize read crossover, we employed 10 bp sequence tags designed to be robust to indel and substitution errors [93]. Further details regarding extraction and preparation protocols per species is described in Table S1. All sequence data have been deposited in the NCBI sequence read archive (SRA).

Reads for all species were trimmed or excluded based on sequence quality scores using Trimmomatic v. 0.36 [94] with default parameters. The Agalma 1.0.1 pipeline [95, 96] was then employed to automate all steps from transcriptome assembly to alignment and construction of data matrices. This phylogenomic workflow allows for straightforward integration of a variety of bioinformatics tasks, including alignment with Bowtie2 [97] and MAFFT [98], assembling with Trinity [99] and alignment trimming with GBlocks [100], among many others (see [95]). Summary statistics output by Agalma for each library are shown in Table S1. Initially, outgroups were represented using transcriptomic datasets, but this resulted in high amounts of missing data. To circumvent this problem, outgroups were replaced with publicly available protein models derived from echinoderm genomes (obtained from [45]). These greatly outperformed transcriptomic data except in the case of the protein model of *Parastichopus parvimensis* H. L. Clark, 1913, which yielded a much lower number of recovered loci than the transcriptome of *Holothuria forskali*. Holothuroids were thus represented using the latter. Given the lack of available protein models for crinoids, all trees were rooted using data derived from the high-quality genome of the hemichordate *Saccoglossus kowalevskii*. The resulting matrix was reduced to a 70% occupancy value, resulting in 1,040 aligned loci.

As already explained, initial analyses were complicated by what we interpret as massive contamination by cidaroid sequences of the transcriptome of *Arbacia punctulata* (Fig. S2). In order to explore the presence of other sources of contamination in the alignment output by Agalma, we used BLAST+ [101] to compare all sequences against a database including protein models for both metazoan and non-metazoan representatives (including common contaminants such as Archaea, Bacteria and Fungi; available at http://ryanlab.whitney.ufl.edu/downloads/alienindex/). For each sequence, the E-value of the best metazoan and non-metazoan hits were used to calculate the alien index (AI), an indicator of foreign (in this case, non-metazoan) origin, as described in [102]. Estimation of AI was automated using alien_index version 3.0 [103]. No sequence in the alignment was shown to have a definite non-metazoan origin (all AI < 45). The analysis was repeated, this time obtaining AIs for a comparison between echinoderm and non-echinoderm metazoans. For this, the protein model for *Strongylocentrotus purpuratus* was removed from the metazoan set and incorporated into a second set including all publicly available protein models for echinoderms (all of the ones used here, plus those of *Parastichopus parvimensis* and *Lytechinus variegatus*). Once again, no echinoderm sequence in our phylogenomic matrix was found to have a definite non-echinoderm origin. Finally, a similar approach to the one used to confirm contamination in *Arbacia* was repeated for all newly-generated transcriptomes. Sequences for each focal transcriptome were compared against two randomly selected transcriptomes using p-distances, and a linear regression was fit to the data. Extreme outliers from this regression line might indicate assembled sequences incorporating foreign reads. Regression residuals are plotted in Fig. S3, showing very few sequences that dramatically deviate from the expected values. In fact, 99.1% of the data fall within a prediction envelope of 3 standard deviations. A wide variety of mechanisms other than contamination can potentially explain the departure of the few remaining sequences from the expected patterns of divergence. Nonetheless, even if these do represent instances of cross-contamination, their effect is not expected to bias systematically the breadth of phylogenetic approaches employed.

ML inference on the concatenated alignment was performed using a variety of approaches to model molecular evolution. Firstly, analyses were run using RAxML-NG v. 0.5.1 [104] on the unpartitioned matrix using the LG4X mixture model, which models heterogeneity across sites using four substitution matrices to which characters are assigned depending on their evolutionary rate [105]. A second mixture model, posterior mean site frequency (PMSF) [106], was explored using IQ-TREE v1.6.6 [107] (-m LG+C60+F+G) as a fast approximation to the CAT family of models. The topology obtained from the ML analysis under the LG4X model was used as guide tree to compute site amino acid profiles. Finally, inference was performed using RAxML v8.2.1 [108] under the best-fit partitioning scheme found using the fast-relaxed clustering algorithm among the top 50% of schemes obtained using IQ-TREE [109] (-m TESTMERGEONLY -mset raxml -rclusterf 50). For this and all other instances of model selection, optimal models were those that minimized the Bayesian Information Criterion. Support was assessed using 200 replicates of non-parametric bootstrapping for the two analyses run in RAxML, and 1,000 replicates of ultrafast bootstrap [110] for the analysis run in IQ-TREE. BI was also performed with the concatenated dataset using two different approaches. In the first, two independent chains of ExaBayes v. 1.5 [111] were run for five million generations using automatic substitution model detection. In the second, PhyloBayes-MPI v. 1.8.1 [112] was run under the site-heterogenous CAT model [113], which models molecular evolution employing site-specific substitution processes. Preliminary runs using the complex CAT+GTR model (two chains, 3,000 generations) failed to converge, as is routinely the case with large phylogenomic datasets [114]. Nonetheless, exploration of the majority rule consensus tree (available online) revealed disagreement among chains regarding a few nodes within camarodonts, with the rest of the topology being identical to that of Fig. 2a. A more thorough analysis was performed under the simpler CAT-Poisson model, with two independent chains being run for 10 million generations. For both BI approaches, stationarity was confirmed using Tracer v1.6 [115], the initial 25% of samples were excluded as burn-in, and convergence was assessed using the software accompanying each program (in both cases, maximum standard deviation of split frequencies = 0).

Finally, coalescent-based species tree inference was performed using the summary algorithm implemented in ASTRAL-II [50], estimating support using local posterior probabilities [116]. Gene trees were estimated in RAxML under the best-fit model for each partition. Species tree reconstruction was then performed using the complete set of 1,040 gene trees, as well as a subset of 354 gene trees obtained after excluding 66% of genes that showed the highest evidence of both among-lineage rate heterogeneity and saturation. Rate heterogeneity was estimated as the variance of root-to-tip distances, and saturation as the slope of the regression of *p*-distances on patristic distances [117]. The value for the slope was subtracted from 1 so that higher numbers correspond to increased saturation. Both metrics were centered, scaled and added together, and genes were excluded if they were among the highest-ranking 66%. Calculations were performed in the R environment [118] with packages *adephylo* [119], *ape* [120], *phangorn* [121] and *phytools* [122] (R code is available at the Dyrad repository). In order to explore topological incongruence, gene trees were decomposed into quartets using SuperQ v. 1.1 [123], and a supernetwork was built in which branch lengths were calculated as the frequency of quartets in the set of ML gene trees using SplitsTree v4.14.6 [124].

To test whether outlier sequences have any impact on the results presented here, we reran a subset of the inference approaches using a further curated dataset. Gene trees were scrutinized using TreeShrink [125], which filtered out sequences with unexpectedly long branches by finding terminals that had strong effects on gene tree diameter (i.e., the maximum distance between pairs of terminals), given species-specific distributions of proportional reduction in gene tree diameter after exclusion. Even though terminal branches can be long for biological reasons, the set of detected outlier sequences was expected to be strongly enriched in erroneous sequences, including those suffering from any lingering issues of contamination, incorrect orthology assessment and misalignment. The algorithm implemented was designed to distinguish between branches that are expectedly long, such as outgroups and fast-evolving species, from branches that are unexpectedly long. However, under default parameters, the set of branches selected as outliers was still strongly enriched in sequences from outgroups and the fast-evolving *Echinocyamus cripsus* (28.0% of outlier sequences compared to 15.6% of representation in the matrix). We therefore reran the software with a reduced tolerance for false positives (-q 0.02), and obtained more robust results. The program suggested the exclusion of 345 sequences, reducing overall occupancy to 69.2%. Analysis of this dataset under coalescent, ML and BI approaches (using ASTRAL-II, IQ-TREE and ExaBayes as already described, see above), revealed no effect of these sequences on the topology, branch lengths or support values (Fig. S4).

We used the Swofford-Olsen-Waddell-Hillis (SOWH) test [51] to evaluate two specific hypotheses of relationships—the monophyly of Acroechinoidea and Clypeasteroida. This topological test compares the difference in log-likelihood scores (*δ*) between the maximum likelihood tree and a constrained topology, obtained by enforcing the monophyly of the clade under consideration, with a distribution of *δ* values obtained via parametric bootstrapping (i.e., using data simulated on the constrained topology). The test was implemented using SOWHAT v0.36 [126], enforcing separate monophyly constraints for Acroechinoidea and Clypeasteroida and setting the model to JTT+Γ+I with empirical amino-acid frequencies (--raxml_model = PROTGAMMAIJTTF), selected as the optimal unpartitioned model by IQ-TREE for the concatenated dataset. Evaluation of the confidence interval around the resulting *P*-values revealed no need for more replicates (i.e., upper limits of the 95% confidence intervals surrounding both *P*-values were < 0.05). Subsequently, log-likelihood scores for all sites in the ML unconstrained and both constrained topologies were obtained using RAxML, allowing gene-wise *δ* values to be calculated (as in [70]). The relationship between these gene-specific *δ* values and several factors with the potential to introduce systematic biases was explored using multiple linear regressions. The predictor variables included the amounts of saturation and branch-length heterogeneity (calculated as explained above), as well as the levels of missing data and compositional heterogeneity. This last was estimated as the relative composition frequency variability (RCFV; [127]) using BaCoCa v1.103 [128]. Only genes that showed some support for either one of the topologies were included in the regression, enforcing an arbitrary cutoff of absolute *δ* values > 3. With this approach, we evaluated both the strength and distribution of signal for our alternative hypotheses, as well as the possibility that this signal is the product of processes other than phylogenetic history.

## Availability of data and materials

The datasets and R code supporting the results of this article will be made available through the Dryad repository. Sequences are deposited in GenBank, and will be made available upon acceptance of the manuscript. Supplementary figures will be hosted on the journal’s website.

## Competing interests

The authors declare that they have no competing interests.

## Funding

This work was supported by a Yale Institute for Biospheric Studies Doctoral Pilot Grant to NMK, US National Science Foundation (NSF OCE-1634172), KAUST, and Scripps Oceanography funds to GWR and a Smithsonian Marine Science Network Grant to SEC. Funds were also contributed by the Yale Peabody Museum Division of Invertebrate Paleontology. The submersible expedition was supported by a Smithsonian “Grand Challenges” grant to HAL, among others.

## Authors’ contributions

NMK and GWR conceived and designed the study. GWR, SEC and HAL sampled many of the specimens used. NMK and SEC performed RNA extractions, prepared libraries and sequenced transcriptomes. NMK performed all data analyses. HAL, DEGB and GWR provided the necessary facilities, as well as assistance at different stages of the study. NMK, RM and GWR interpreted the results, and NMK and RM wrote the manuscript with input from all other authors.

## Acknowledgments

Many thanks to Tim Ravasi for providing resources and collection facilities to GWR via King Abdullah University of Science and Technology (KAUST). We thank Chief Scientist Erik Cordes, the captain and crew of the RV *Atlantis* and the crew of the HOV *Alvin* for assistance in specimen collection off Costa Rica. Thanks also to David Clague (Monterey Bay Aquarium and Research Institute) for inviting GWR to Juan de Fuca and to the captain and crew of the RV *Western Flyer* and the crew of the ROV Doc Ricketts for specimen collection. The authors are also grateful to Gustav Paulay, Phil Zerofski, Fredrik Pleijel, Frédéric Ducarme and Alejandra Melo for their sampling efforts; Josefin Stiller, Ekin Tilic and Avery Hiley for their help with molecular work; Charlotte Seid for the handling and cataloging of specimens at Scripps; and Casey Dunn, Kaylea Nelson and Benjamin Evans for computational assistance. Specimens of *Mellita tenuis* were purchased from Gulf Specimen Marine Laboratory. *Conolampas sigsbei* was collected in the submersible “Curasub” on time donated by Adrian “Dutch” Schrier and identified by David L. Pawson.

